# Times birds need to evolve mating cues under allopatry and parapatry

**DOI:** 10.1101/2024.03.01.582980

**Authors:** Richard M. Sibly, Robert N. Curnow

**Author notes:** Corresponding author. Mailing address School of Biological Sciences, University of Reading, Reading RG66EX UK.

## Abstract

The time needed for the evolution of mating cues that distinguish species, such as species-specific songs or plumage coloration, has received little attention. Motivated by observations of rapid divergence in mating cues in birds we here compare speciation times under allopatry with those under parapatry. We assume that under allopatry the mating cues are neutral but under parapatry may be acted on by selection. The expected time needed for a neutral substitution at a mating-cue locus is approximately equal to the reciprocal of mutation rate, but the time needed under parapatry can be much shorter. The time needed is reduced by a factor of 4 x (population size) x (selection coefficient on the mating cue). In the simplest case of parapatric speciation the mating cue is an adaptive magic trait such as beak size which is acted on by selection directly, but this does not affect traits like species-specific songs or plumage colouration that are not acted on directly by selection. In an alternative model of parapatric speciation in which the locus coding for a neutral mating cue is not linked to loci producing local adaptation, local adaptation induces selection on the mating cue. We show quantitatively how, with complete phenotype matching, speciation times can be obtained from population size, migration rates between subpopulations and the ecological selective conditions within them. Our results suggest that only parapatric, not allopatric, speciation can account for the times recorded for the evolution of species-specific songs or plumage coloration.

## Introduction

Speciation in birds – and many other species – is currently considered generally to occur after a period of allopatry, during which a population becomes split into geographically isolated subpopulations which remain separate and diverge over thousands of generations, and eventually can no longer interbreed (Tobias et al., 2020). Allopatric speciation is not the only possible speciation mechanism, however, and in recent decades attention has been given to the role of divergent natural selection, which leads to local adaptation and, potentially, speciation in the presence of gene flow (see reviews in, e.g., (Janicke et al., 2019; Kopp et al., 2018)). Here in addition to allopatric speciation we consider in particular the case of parapatric speciation. The difference between allopatric and parapatric speciation is that there is no migration between subpopulations in the former whereas the latter has migration between regions that experience different ecological selection pressures. Parapatric speciation is thought to be more likely under phenotype matching than under alternative methods of mate choice (Kopp et al., 2018). Phenotype matching means that individuals with a particular mating cue tend to mate with others with the same mating cue. Here we assume that phenotype matching is complete, so individuals only mate with others with the same mating cue. For certain combinations of migration rates and selection coefficients, with phenotype matching parapatric speciation occurs readily in locally adapted populations (Felsenstein, 1981; Sibly and Curnow, 2022).

We consider here two types of parapatric speciation. In the first, which we designate Type 1 parapatric speciation, adaptive traits are used as mating cues, i.e., are ‘magic traits’. The term magic trait refers here to an adaptive trait that is used as a mating cue (see, e.g., (Kopp et al., 2018)). An example is choosing mates on the basis of having a beak size that is advantageous in the niche in which the population lives. The combination of phenotype matching and a magic trait is known to be particularly favourable for speciation (Smadja and Butlin, 2011). Treating a related case, (Servedio and Burger, 2020) used a deterministic haploid model to analyse what they term ‘pseudomagic traits’, in which a mating cue locus and an ecological trait locus are separate but linked. (Servedio and Burger, 2020) show that evolutionary outcomes are similar to Type 1 if the ecological trait locus is tightly linked to the mating cue locus.

Alternatively (Sibly and Curnow, 2022) used a deterministic diploid model to show quantitatively how local adaptation induces selection on mating cues for the case that a locus controlling mating cues is not linked to a locus controlling local adaptation. We refer to this as Type 2 parapatric speciation. Speciation occurs in (Sibly and Curnow, 2022)’s model because individuals in each niche avoid mating with incomers who predominantly carry disadvantageous alleles. The model incorporates a (Felsenstein, 1981) “two-allele mechanism” (see also (Butlin et al., 2021)). Whether the population will speciate depends in (Sibly and Curnow, 2022)’s model as in Felsenstein’s on the balance between migration and selection: some selection at the ecological locus is necessary. (Sibly and Curnow, 2022)’s results were based on simulations that assumed complete phenotype matching, an infinite population divided equally between two niches, migration rates the same in both directions, and local adaptation the result of two alleles P and Q at a single diploid locus. The mating cue is controlled by a single locus with only D alleles prior to a new mating cue produced by a C allele arising by mutation. C is assumed dominant to D. We retain this notation below.

Our focus here is mainly on birds, in which speciation has been extensively studied (Price, 2008). Bird species are generally distinguished by the colour plumage and/or song used in mate choice. Novel mating cues that distinguish species evolve during the process of speciation but the time needed for the evolution of novel mating cues, the subject of the present paper, has received little attention. Analysis of DNA sequences shows that in some cases new mating cues have evolved rapidly. Noteworthy examples occur in the tanagers including southern capuchinos seedeaters (*Sporophila*) and Darwin’s finches. The Neotropical southern capuchinos radiated within the last one million years to form 10 predominantly sympatric species that differ primarily in male plumage coloration and song (Campagna et al., 2017; Campagna et al., 2012; Turbek et al., 2021). Eight of these species emerged in less than 50,000 generations (Hejase et al., 2020). In Darwin’s finches, radiations of ground and tree finches began around 100,000– 300,000 years ago (Lamichhaney et al., 2015). Motivated by these observations of rapid divergence in mating cues in birds, the purpose of this paper is to estimate the time to fixation of a neutral substitution for a novel mating cue under models of allopatric and parapatric speciation. The estimated times to fixation can then be compared with the above data. Although waiting times to speciation have been considered previously this has been mainly in models where incompatibilities accumulate or populations respond to divergent selection (see, e.g. (Gavrilets, 2014)). In this paper the focus is on mating isolation and so the question addressed here is quite new.

Bird speciation is used as a key example though the results have wider application. Although avian coloration genetics has shown that many genes may be involved in plumage coloration (Price-Waldman and Stoddard, 2021), the number of possible mutations that could result in phenotypic change is not known. Our initial analysis is therefore based on calculating times at a single given locus, but times are reduced if there are several loci that can result in phenotypic change, as described in the Discussion.

The paper is in two sections. In the first we show that the time needed for a new neutral mating cue to evolve is much longer under allopatry than under parapatry. In the second section we calculate for selected cases the time needed under Type 2 parapatry, in which local adaptation induces selection on mating cues even though the loci controlling the mating cues are not close to the loci controlling local adaptation. This second section offers some insight into how the time needed under Type 2 parapatry depends on the selection coefficients in the niches containing the parapatric population. We hope that our main results, showing quantitatively how parapatry reduces speciation times, will be of interest and provide a starting point for further work relaxing model assumptions and studying different models for mate choice.

### Section 1: Time for a new mating cue to spread to fixation

In this section we compare times for a new mating cue to arise and spread to fixation under allopatry and parapatry.

## Methods

We assume mating cues are controlled by a single locus which is homozygous prior to a new mating cue arising by mutation. The alleles at this locus are labelled D alleles. Any mutation of D can produce a new mating cue. We label C the first such mutation that spreads to fixation, and assume C dominant to D. In Type 1 parapatric speciation, the mating cue is an adaptive trait and so directly subject to natural selection. In allopatry or Type 2 parapatry, mating cues are not subject to selection directly, i.e. they are selectively neutral, but under Type 2 parapatry they may be acted on by selection indirectly as a result of phenotype matching (Sibly and Curnow, 2022). We assume that all individuals irrespective of frequency of their mating cue do mate. Our aim in this section is to use standard methods of population genetics to calculate the times for new mating cues to arise and spread to fixation as a result of the mating cue selection coefficient acting directly or indirectly on C. This selection coefficient is assumed to be zero under allopatry, but positive under parapatry.

The time of arising of the first mutated allele C that goes to fixation, here labelled T_a_, and the time from this mutation arising to fixation, T_f_, are random variables, so the expected time to the fixation of C, E[T], can be calculated from:

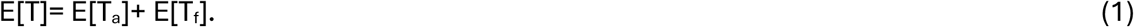

The time of arising of the first mutated allele that goes to fixation depends on the mutation rate. We let µ be the rate at which new neutral mutations arise per allele per generation. We begin by calculating the expected time needed to acquire a new C mating cue under allopatry.

### Timings under allopatry

Under allopatry we assume that mating cues are selectively neutral. To calculate the expected time of arising of the first C mutation that goes to fixation, E[T_a_], we begin by calculating the probability of a neutral mutation arising and going to fixation. This is the product of the probability of a mutation arising, µ, and the probability it goes to fixation. Assuming large N and small *μ*, the per generation probability of a mutation arising is approximately 2*Nμ*, where *N* is the number of individuals in the population. This is because there are 2N alleles at the focal locus and the chance of each mutating is *μ*.

The probability the arising mutation goes to fixation is 1/(2N) if the mutation is neutral ((Kimura and Ohta, 1971), see also (Otto and Whitlock, 2013)). So the probability of a neutral mutation arising and going to fixation is 2*Nμ*/(2*N*) = *μ*, and the expected time of arising of mutations that go to fixation is the reciprocal of this, 1/*μ*.

Based on diffusion approximations with large N and continuous time, the expected time from the arising of a neutral mutation until it reaches fixation is approximately 4N generations, conditional on the allele fixing ((Kimura and Ohta, 1971), see also (Otto and Whitlock, 2013)). The expected time to the fixation of C is now given by eqn (1) as:

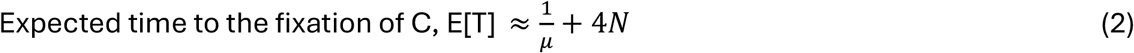

### Timings under parapatry

We consider here two types of parapatric speciation, both of which assume the existence of phenotype matching and migration between subpopulations which differ in ecological selection pressures. In Type 1 parapatric speciation, the novel mating cue C is adaptive, i.e., it codes for a ‘magic trait’. In this case completion of local adaptation and speciation coincide. In Type 2 parapatric speciation, local adaptation induces selection on a mating cue that is not linked to an adaptive trait. The strength of selection on the mating cue then depends on how selection operates on the adaptive trait. The overall selection coefficient on the mating cue depends on the selection coefficients operating on the adaptive trait – positive in one niche and negative in the other. The relationship between these selection coefficients is described in detail for Type 2 parapatry in Section 2, below. In considering the timing of parapatric speciation, what is important is the overall selection pressure on the trait under consideration, which could be either the adaptive trait or a mating cue distinct from the adaptive trait. The overall selection pressure on the trait can be quantified by an *overall selection coefficient*, here designated *s*. In this section s is taken to be constant. This is approximately true in Type 2 parapatry except for the first few generations of the spread of the mating cue, when it is higher (example in Fig. 2b, below).

We now use equation 1 to calculate the expected time to the fixation of C, E[T]. The probability of a mutation arising is approximately 2*Nμ*, as before. The probability the arising mutation goes to fixation is approximately 2*s* if *s* is small ((Kimura and Ohta, 1971), see also (Otto and Whitlock, 2013)). So the probability of a selected mutation arising and going to fixation is 4*Nμs*, and the expected time of arising of selected mutations that go to fixation, T_a_, is the reciprocal of this, 1/(4N*μs*).

The time from C arising to fixation can be obtained from equations given by (Charlesworth, 2020). These equations apply to cases where the fixation index, F, is constant and known. Under parapatry, however, as C spreads, mating is increasingly non-random and F increases. This situation has not been modelled, and we therefore consider two limiting cases: F = 0 (random mating), and F positive and constant. It turns out that the time from C arising to fixation is longer for random than for non-random (F > 0) mating, so random-mating times represent a limiting case – actual times will be shorter than those for random mating.

With random mating, for a dominant allele at an autosomal locus with weak selection and a large population, (Charlesworth, 2020) showed that the time from C arising to fixation in units of 2N, conditional on the allele fixing, is approximately:

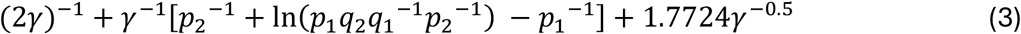

where *γ* = 2*Ns*, assumed >>1, *p*_1_ = 1 − (2*γ*)^−1^, *p*_2_ = 0.8862*γ*^−0.5^, q_1_=1-p_1_, q_2_=1-p_2_, and N_e_ is taken to be N.

With assortative mating and fixation index F, (Charlesworth, 2020) showed that the time is approximately:

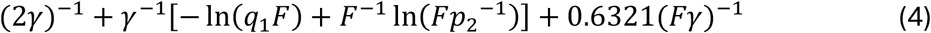

In deriving eqn 4 it is assumed that *F* > *O*(*s*), and *p*_2_^−1^is large. Numerical evaluations of eqns 3 and 4 reported in Supplementary Material 1 show that times from C arising to fixation are always lower without random mating than with it; the ratio declines as F or Ns increase, getting below 5% when Ns>100,000 and F>0.1. The expected time to the fixation of a selected mutation for the limiting case of random mating can now be approximated from equation 1 as:

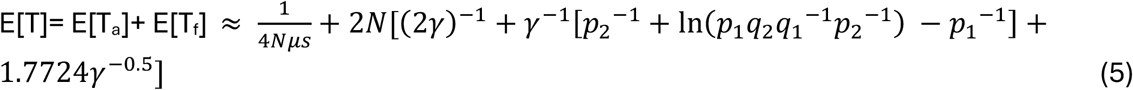

Equations 3 – 5 hold provided that s < 0.02 and Ns is large (Charlesworth, 2020).

## Results

The expected times to fixation of a mating cue for allopatry and parapatry are shown in relation to the selection coefficient *s* in Fig. 1, for the limiting case of random mating. Both T and T_a_ are plotted but in Fig. 1a these overlay if N < 100,000 indicating that time from C arising to fixation is then negligible in comparison with the time of arising of the first mutated allele that goes to fixation, T_a_. Fig. 1a shows that mating cues generally evolve substantially faster under parapatry than under allopatry. With µ=10^−8^, *s* = 0.02 and a population size of 10,000, cues evolve in 100,000 generations under parapatry, compared with 100 million generations needed under allopatry. In a population of 100,000, cues evolve in a few thousand generations under parapatry. Even for a population of 100 individuals, cues evolve faster under parapatry than allopatry provided *s* > 0.03.

**Fig. 1.**
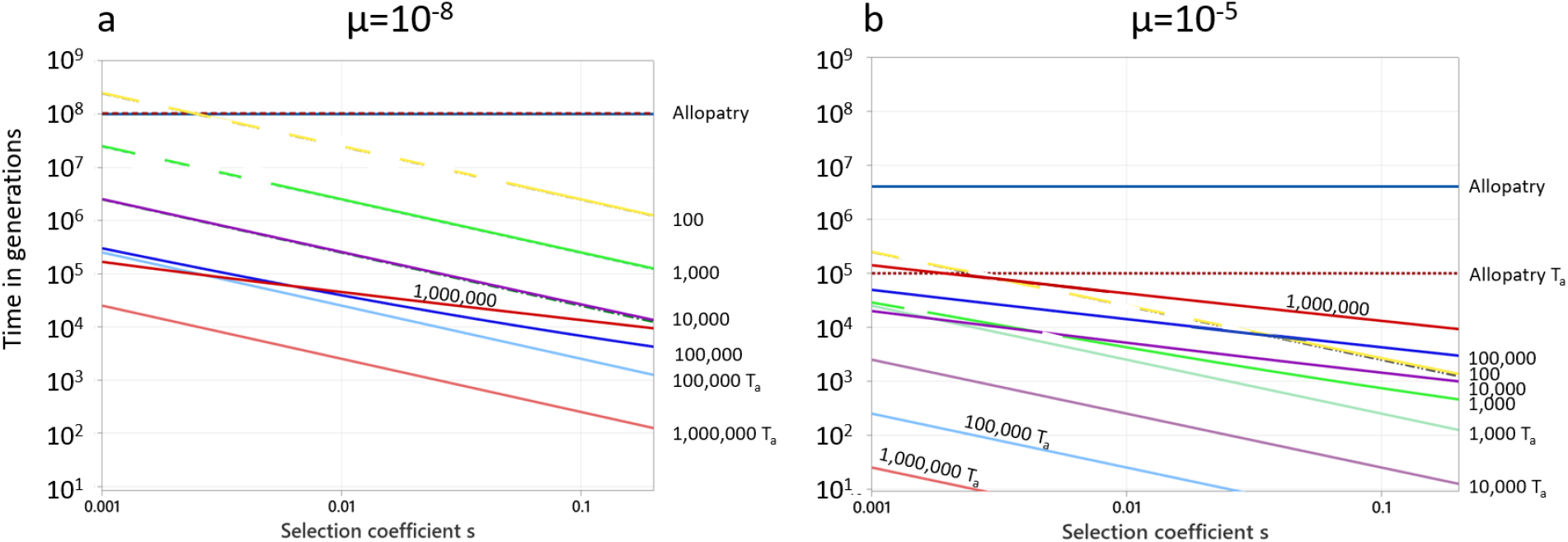
Expected times to fixation of a mating cue under allopatry and parapatry for the limiting case of random mating. Times to fixation are plotted against the overall selection coefficient *s* on log-log scales for two mutation rates: a) µ=10^−8^; b) µ=10^−5^. The lines for allopatry are shown for comparison with those for parapatry but the selection coefficient under allopatry is assumed to be zero. The allopatry lines satisfy equation 2. The other lines are for parapatric populations, for which the numbers in the right-hand columns indicate population size in equation 5. The lines for both E[T] and E[T_a_] are plotted and these generally overlay. Where they do not overlay the E[T_a_] line has the suffix T_a_, the E[T] line has no suffix. The cases in which the lines overlay are in a) where N < 100,000 and allopatry; and in b) where N < 1,000. Dashed lines indicate regions where Ns <5, i.e., failure of the assumption Ns large that is used to derive eqns 3 – 5.

**Fig. 2.**
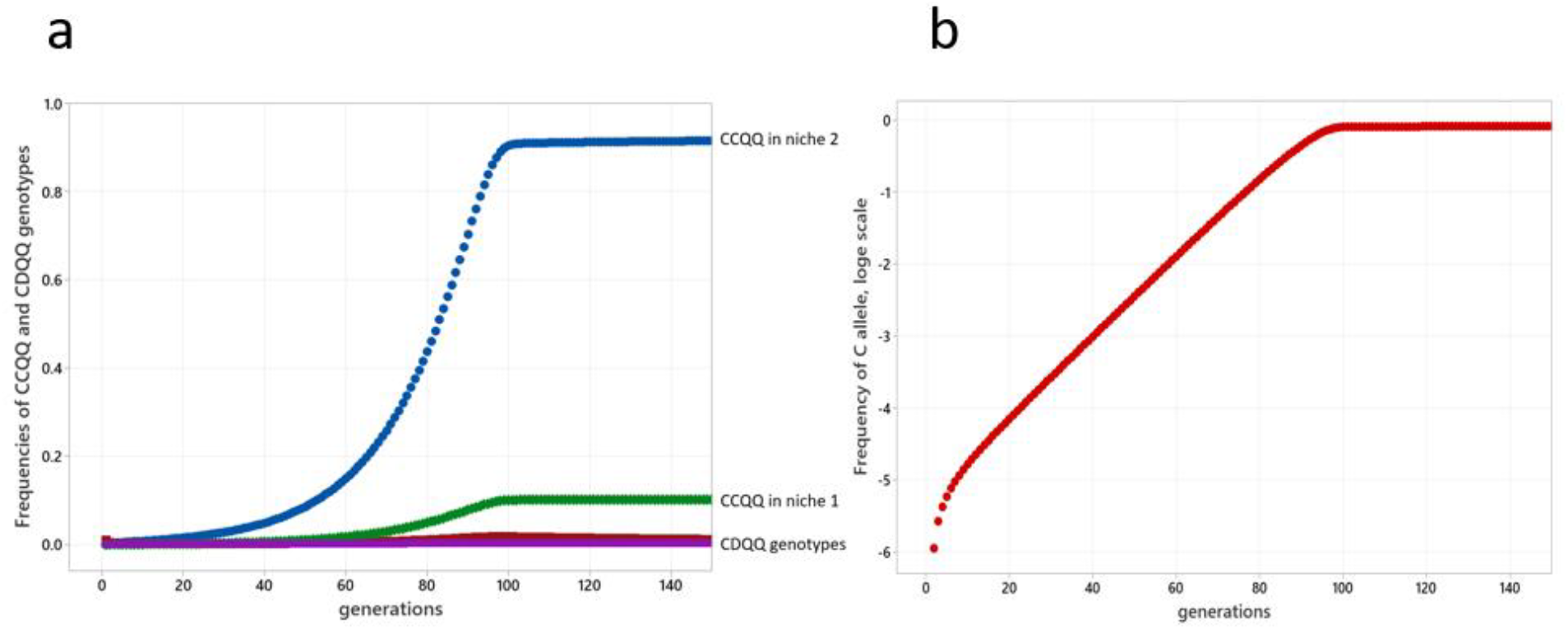
Example of the exponential increase of the C allele in the (Sibly and Curnow, 2022) model. a) The frequency of the CCQQ and CDQQ genotypes in the two niches. The C allele was originally introduced at low frequency (1%) into niche 2 as a CDQQ genotype. The evolutionary outcome is that only two genotypes persist in the two niches, CCQQ and DDPP (DDPP is not shown here). b) The resulting increase of the C allele is approximately linear on a log scale, indicating that the overall selection coefficient s is constant except for the first few generations, when it is higher. Parameter values were m=0.067, s_2_ = 2.0, s_1_ = -0.4. Simulation carried out as in (Sibly and Curnow, 2022).

Fig. 1a shows the situation when the mutation rate µ=10^−8^. If the mutation rate is higher, time to fixation is reduced, and the contribution of the time from C arising to fixation is more pronounced. These effects are illustrated in Fig. 1b, where µ=10^−5^. The colour coding is the same in Figs. 1a and b. All times are reduced in Fig. 1b in comparison with Fig. 1a. Times to fixation under parapatry are closer together than in Fig. 1a because of increases in the contribution of the time from C arising to fixation: these increase with N. Over the ranges of s and N shown in Fig. 1b, times to fixation range from under a thousand if s=0.2 and N=1,000, to a little over 100,000 if s= 0.001 and N=1,000,000. These are much less than the 5 million generations needed under allopatry.

The results shown in Fig. 1 show the effects of selection on the time needed for a trait to evolve. The case labelled ‘Allopatry’ corresponds to the trait being neutral. This is the limiting case as the selection coefficient *s* tends to zero. The cases when *s* > 0 are considered as cases of parapatry. Although these results apply to any evolving traits, our focus here is on discussing them in relation to allopatry and parapatry. Fig. 1 suggests that the time needed for a new mating cue to evolve can be much shorter under parapatry than under allopatry. If the mutation rate µ=10^−8^ per site per generation, the value commonly reported in reptiles, birds, fish and mammals (Bergeron et al., 2023), then in populations under 100,000 the time needed for a new mating cue to evolve depends almost entirely on the time of arising of the first mutated allele C that goes to fixation, as shown in Fig. 1a. The times needed for a new mating cue to evolve under allopatry and parapatry then simplify to 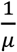 and 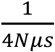 respectively. Thus, the time needed under parapatry is lower than that needed under allopatry by a factor of 4*Ns*. On the log-log scales of Fig. 1, the relationship for allopatry is log(*time needed*) = − log *μ*. The relationship for parapatry is log(*time needed*) = − log *μ* − log 4 − log *N* − log *s*. So, *time needed* can be decreased by an order of magnitude by increasing population size *N* by an order of magnitude, or increasing the overall selection coefficient *s* by an order of magnitude.

If the mutation rate is higher, time to fixation is reduced, and the contribution of the time from C arising to fixation is more pronounced (example in Fig. 1b). The net effect is that the time needed for a new mating cue to evolve is still much shorter under parapatry than under allopatry. Allowing for changes in the y-axis scale, results are similar to those shown in Fig. 1 as mutation rate is varied from µ=10^−8^ to µ=10^−2^ (results not shown).

In this section we have shown that the time needed for a new mating cue to evolve can be much shorter under parapatry than under allopatry, as illustrated in Fig. 1. This is true of both types of parapatric speciation.

### Section 2: Dependence of the mating cue selection coefficient on the selective regimes in two niches in (Sibly and Curnow, 2022)’s Type 2 parapatric populations

The objective in this section is to calculate the time needed for a new mating cue to evolve when local adaptation induces selection on mating cues even though the loci controlling the mating cues are not linked to the loci controlling local adaptation – i.e., Type 2 parapatry. The mating cue is assumed to be coded by a C allele arising by mutation at a locus not linked to another locus which produces local adaptation.

Without local adaptation there would be no selection at the mating cue locus, but in the presence of local adaptation selection occurs at the mating cue locus. We term the resulting selection coefficient the overall selection coefficient on the mating cue. In this section we consider how the overall selection coefficient on the mating cue relates to the selection coefficients operating on the adaptive trait – positive in one niche and negative in at least one other. An indication of the answer to this question can be obtained using (Sibly and Curnow, 2022)’s two-niche model of Type 2 parapatric speciation.

## Methods

In (Sibly and Curnow, 2022)’s model generations are discrete and individuals die after mating. At the start of each generation individuals in each niche mate with others of the same mating-cue phenotype, and all mating individuals obtain the same number of offspring. The number of offspring of each genotype that survive in each niche is the product of its initial frequency and its fitness. Population regulation then returns population numbers to their initial values, after which some individuals migrate between niches. Local adaptation is the result of two alleles P and Q at a single locus. PP has fitness 1 in both niches. QQ has fitness 1+s_1_ in niche 1 and 1+s_2_ in niche 2. s_2_ is assumed positive and s_1_ negative. Whether or not local adaptation occurs depends not only on the values of s_1_ and s_2_ and the level of dominance of Q, but also on the migration rates between the two niches. Further details of the model including the recursion equations showing how genotype frequencies change between generations are given in Supplementary Material 2.

For the case that population sizes in the two niches are equal and migration rates in both directions are the same, (Sibly and Curnow, 2022) used deterministic simulation of the evolutionary process to map the set of migration and selection rates for which local adaptation occurs, and showed that in all analysed cases, the evolutionary outcome is speciation if a C allele coding for a new neutral mating cue arises by mutation. C allele was originally introduced into the simulation at low frequency (0.05%). As the C allele spread the CD and PQ heterozygotes were eliminated, so that eventually the population consisted only of the CCQQ and DDPP genotypes. The C allele increased in frequency approximately exponentially until close to its final value (example in Fig. 2). During the exponential phase the mating cue selection coefficient on C alleles was obtained by simulations carried out as in (Sibly and Curnow, 2022). By repeating this procedure over a range of values of s_1_, s_2_ and migration rate *m*, a picture was built up of the dependence of s on s_1_, s_2_ and m.

## Results

Fig. 3 shows how the mating cue selection coefficient on the C allele, *s*, is affected by migration rate *m* and the selection coefficient *s*_*2*_ in niche 2, for two values of s_1_. s_2_ and s_1_ represent the strength of selection for and against the Q allele in niches 2 and 1 respectively. Note first that selection for C, i.e. s > 0, only occurs within a restricted region in the m s2 plane. This is the region in which local adaptation at the PQ locus is possible. Outside this region C is not selected or is selected against, so s ≤ 0. The highest values of the selection coefficient are 0.013 in Fig. 3a and 0.12 in Fig. 3b. They occur on horizontal ridges where the opposing selection conditions in the two niches are close to being equal and opposite. Thus, the ridge occurs at s_2_ ∼ 0.05 in Fig. 3a and at s_2_ ∼ 0.40 in Fig. 3b. High values of s also occur on vertical ridges at m ∼ 0.05 in Fig. 3a and m ∼ 0.15 in Fig. 3b. Either side of the ridges, s declines down to zero at the edges of the region in which local adaptation is possible. To the left s declines to zero when m = m = 0 corresponds to allopatry since there is no migration between niches. Towards the bottom of Fig. 3, s declines steeply and becomes negative as s_2_ declines to zero. When s_2_ is zero, s is negative because there is no selection for CCQQ in niche 2 and selection against CCQQ in niche 1. s is zero at the upper right of the figure, because local adaptation is not possible there: the region consists entirely of Q alleles, and there is then no selection for C alleles.

**Fig. 3.**
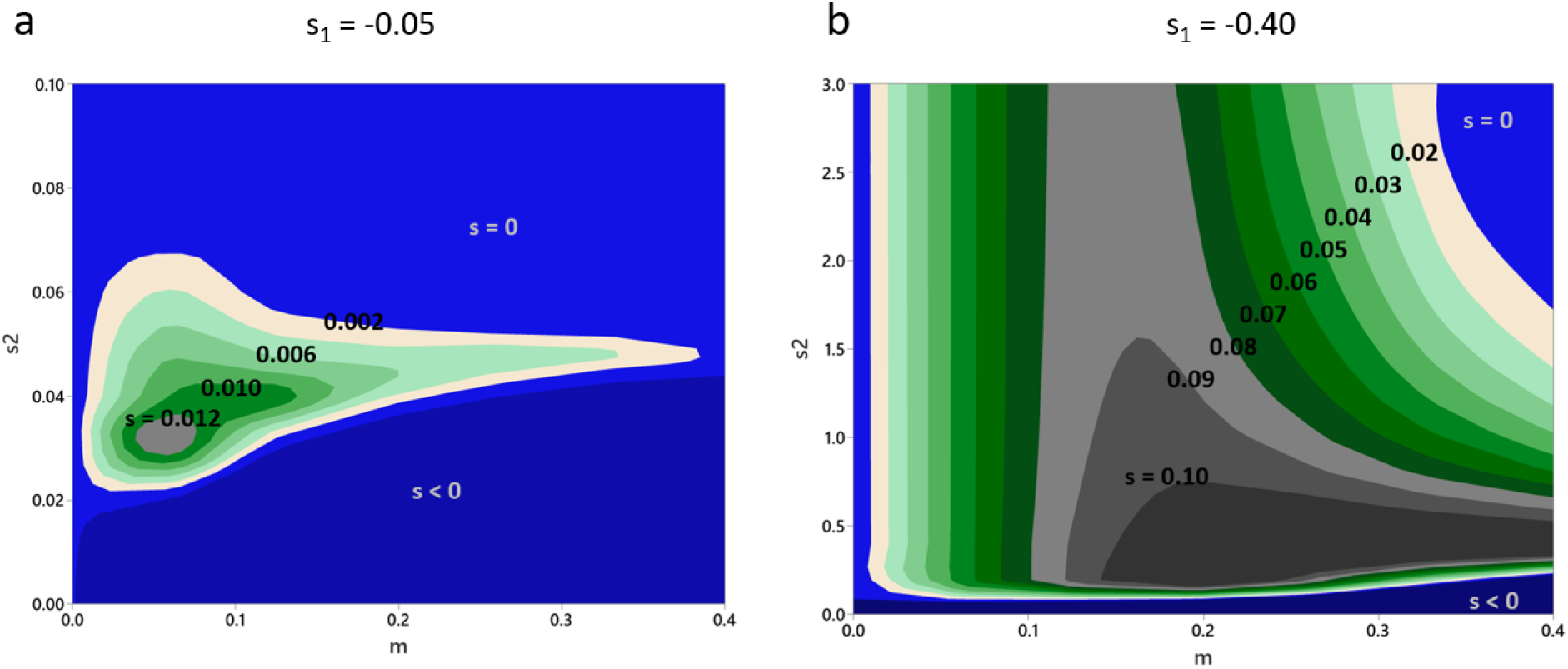
3D contour plots showing how the mating cue selection coefficient, *s*, is affected by migration rate *m* and *s*_*2*_, for two values of s_1_. Mating cue selection coefficients *s* were calculated over grids of points in (Sibly and Curnow, 2022)’s Fig. 2 as follows: (a) corresponds to (Sibly and Curnow, 2022)’s bottom left panel; (b) to their bottom right panel. In these simulations Q was dominant to P. Simulations carried out as in (Sibly and Curnow, 2022). *s* was calculated in the phase of exponential increase of C, generation 50 in (a) and 20 in (b). The contour plots were obtained by interpolation between grid points using Minitab 21.

In sum, as in (Sibly and Curnow, 2022) there is no selection for the mating cue outside the region is which local adaptation occurs. Within the region of local adaptation, selection on the mating cue seemingly increases with migration rate from zero up to some maximum, and then declines back down to zero at the edge of the adaptive region. Mating cue selection is also relatively high if the opposing selection conditions in the two niches are approximately equal and opposite, i.e., s_2_ ∼ - s_1_.

The cases analysed assume complete phenotype matching, populations divided equally between two niches, migration rates the same in both directions, and local adaptation the result of two alleles P and Q at a single locus. The mating cue selection coefficient s is at most 0.12 in the cases analysed in Fig. 3, within the range of validity of the assumptions used to derive eqns 3 - 5. Complete phenotypic matching is assumed in Fig. 3, but if some individuals mate with others of a different phenotype then the mating cue selection coefficient would be reduced. The evolution of choosiness could be explored using the methods and results of (Aubier et al., 2023) as a starting point, and theoretical equations with which to investigate this and other parameter variants are available in (Sibly and Curnow, 2022), though further work is needed to see what happens when key genes are not dominant.

## Discussion

Our main results, illustrated in Fig. 1, show quantitatively how the time needed for a mating cue to evolve depends on the strength of selection for the mating cue. These results hold for any newly evolving allele but our focus in this paper is on mating cues. The results demonstrate quantitatively how much shorter is the time needed for a new mating cue to evolve under parapatry compared with under allopatry. The time needed can be reduced by several orders of magnitude, depending on population sizes and the strength of selection, as shown in Fig. 1. How the strength of selection for the mating cue depends on the strength of selection for an ecological trait in (Sibly and Curnow, 2022)’s Type 2 parapatry is addressed in Fig. 3. To see the relationship between Figs. 1 and 3, consider a fictitious example: suppose an F_ST_ value of 0.31 is recorded for a locus critical to local adaptation, the genotypes varying between two niches. Using the results of (Sibly and Curnow, 2023), the selection coefficients on one of the alleles would be 0.4 in one niche and -0.4 in the other if the absolute values of the selection coefficients were equal. Fig. 3 shows that the mating cue selection coefficient would then be around if the migration rate was 0.15, in which case from Fig. 1 the expected speciation time would be about 26,000 generations if the population size was 100,000. For smaller populations or lower migration rates the expected speciation time would be longer. So from measured F_ST_ values for an ecological trait, if we have some indication of migration rates we can get an idea of expected speciation times in (Sibly and Curnow, 2022)’s Type 2 parapatry.

These results assume that phenotypic change can only occur as a result of genetic change at a single locus. If there are several loci that potentially could code for a new mating cue, the number being given the symbol *x*, then *μ* should be replaced by *μx* throughout the analysis. This is because there are now 2*Nx* alleles at the *x* focal loci and the chance of each mutating is *μ*, so the per generation probability of a mutation arising is approximately 2*Nμx*. Considering only the first mutation that goes to fixation, the rest of the analyses follow as before with *μ* replaced by *μx*. The effect is identical to that of increasing *μ*, so that for example in comparison with Fig. 1a, Fig. 1b could result either from increasing *μ* by three orders of magnitude or from an equivalent increase in *x*. So speciation times are reduced the more loci there are that potentially could code for mating cues.

The conditions under which speciation occurs are the same in Types 1 and 2 parapatry. This is because in both cases speciation occurs only if the ecological selection pressures are such as to produce local adaptation. This follows for Type 1 from the way it is defined, for Type 2 see (Sibly and Curnow, 2022). If the mating cue evolves by Type 1 parapatric speciation, the mating cue is the same as the adaptive trait – a prime example is beak size. Speciation would be rapid, but would not involve evolution of species-specific songs or plumage coloration unless these were themselves locally adaptive. By contrast species-specific songs or plumage colorations do evolve under Type 2 parapatric speciation, induced by the ecological selection pressures that produce local adaptation.

The assumptions and approximations used in deriving equations 2 – 5 merit discussion. It has been assumed that mating cues under allopatry are selectively neutral, but this might not be true if sexual selection operated on the mating cues during allopatry.

However to get differences between the geographic regions it would be necessary for sexual selection to operate differently in the different regions. Sexual selection depends on mate-choosers – often females – having preferences for selected traits, so these preferences also would need to differ between the geographic regions. So while possible in some cases this seems unlikely as a general explanation, and we contend that our assumption of neutrality of mating cues under allopatry is an appropriate theoretical starting point. It is also assumed that assortative mating has no costs, but being choosy may reduce the chance of mating, reducing the mating cue selection coefficient in the parapatric models (Aubier et al., 2023; Kopp and Hermisson, 2008; Schneider and Burger, 2006). The probability of a mutation arising is 1 − (1 − *μ*)^2*N*^ and this is approximated as 2*Nμ*. This is a good approximation even when *N*=1,000,000: with *μ* =10^−8^ the approximation then gives 0.0200 compared to the exact value of 0.0198. In deriving equations 3 – 5, s is taken to be constant, though in reality it is higher in (Sibly and Curnow, 2022)’s Type 2 parapatry in the first few generations of the spread of the mating cue, as shown in Fig. 2b. The effect of this will be to shorten the time to fixation of a mating cue under Type 2 parapatry. Fig. 1 only shows s values up to 0.2 and Ns large, the regions in which (Charlesworth, 2020)’s eqns 3 – 5 are valid. In sum, since the values of *s* may often be less than 0.1 (Fig. 3), our conclusions are unlikely to be affected by the approximations used in deriving equations 2 – 5.

We are now in a position to compare with data the expected times to fixation of a mating cue under allopatry and parapatry shown in Fig. 1. As described in the Introduction, analysis of molecular phylogenies suggest that eight species of southern capuchinos seedeaters (*Sporophila*) evolved distinct male plumage coloration and in some cases song in less than 50,000 generations, and in Darwin’s finches radiations of ground and tree finches began around 100,000–300,000 years ago and at least in some cases involved species-specific songs (Grant and Grant, 2018). These timings of around 100,000 generations are three orders of magnitude less that the time of 10^8^ generations that Fig. 1 suggests would be required under allopatry, but are similar to those resulting from Type 2 parapatry. Although there are several simplifications in our argument, it seems likely that only Type 2 parapatric speciation can account for the times recorded for the evolution of species-specific songs or plumage coloration.

Our results encourage further research on species that are in the process of speciating or have very recently speciated. A key question is how phenotype matching is achieved. A likely candidate in birds is sexual imprinting, a process whereby individuals choose mates that resemble other individuals, usually one of their parents. Sexual imprinting seems to be a general feature of birds, shown to exist in over 100 species belonging to 15 different orders, and in both sexes (ten Cate and Vos, 1999), and has also been found in mammals, fish (cichlids and stickleback) and frogs (Verzijden et al., 2012; Yang et al., 2019). Phenotype matching could alternatively be achieved if the gene(s) determining mating cues were either identical to or close to loci determining mate preferences. Comparing the genomes of very recently diverged species, under Type 2 parapatry we expect to find divergence at just one plumage/song locus, since the evolution of more loci, while they may clarify the signal, are likely to add substantially to the time taken. In addition to the divergence at the plumage/song locus we expect to find divergence at one or more loci conferring local adaptation(s), which we expect to have diverged earlier, on the premise that, under Type 2 parapatry, local adaptation is in general a necessary precursor to the evolution of a new mating cue.

Finally we offer the thought that with phenotype matching of mating cues, parapatric speciation is a means of enhancing local adaptation. Although our modelling assumptions are restrictive so that caution is needed in comparing the results to empirical data, we hope that our main results, showing quantitatively how parapatry can reduce speciation times, will be of interest and provide a starting point for further work relaxing model assumptions or studying different models for mate choice.

## Acknowledgements

We are very grateful to Roger Butlin and Brian Charlesworth for comments on the manuscript.

## Conflict of Interest

None.

Supplementary Material 1: Numerical evaluation of times to fixation given by equations 3 and 4.

Supplementary Material 2: Extended methods for Section 2.

