## Supplementary Material 1 for "Times birds need to evolve mating cues under allopatry and parapatry"

**Supplementary Material 1: Numerical evaluation of times to fixation given by equations 3 and 4**

Table S1. Numerical evaluation of times to fixation given by equations 3 and 4 for selected cases satisfying Ns≥1000, s<0.02 and p_2_^-1^≥50. These times are in units of 2N. The last column shows the ratio of the time from equation 3, to that from equation 4. These ratios are plotted against log_10_(Ns) in Fig. S1.

| log10 (Ns) | log10 (F) | gamma | F | p1 | p2 | time to fixation eqn 3 | time to fixation eqn 4 | ratio eqn 4 to eqn 3 |
| --- | --- | --- | --- | --- | --- | --- | --- | --- |
| 3 | -2 | 2000 | 0.01 | 0.99975 | 0.0198 | 0.070712 | 0.00411 | 0.058113 |
| 4 | -2 | 20000 | 0.01 | 0.99998 | 0.0063 | 0.02127 | 0.00628 | 0.29537 |
| 5 | -2 | 200000 | 0.01 | 1 | 0.002 | 0.00658 | 0.00122 | 0.184726 |
| 6 | -2 | 2000000 | 0.01 | 1 | 0.0006 | 0.002062 | 0.00018 | 0.087409 |
| 7 | -2 | 20000000 | 0.01 | 1 | 0.0002 | 0.00065 | 2.4E-05 | 0.03677 |
| 8 | -2 | 200000000 | 0.01 | 1 | 6E-05 | 0.000205 | 3E-06 | 0.014503 |
| 3 | -1.5 | 2000 | 0.031623 | 0.99975 | 0.0198 | 0.070712 | 0.02351 | 0.332454 |
| 4 | -1.5 | 20000 | 0.031623 | 0.99998 | 0.0063 | 0.02127 | 0.00429 | 0.20152 |
| 5 | -1.5 | 200000 | 0.031623 | 1 | 0.002 | 0.00658 | 0.00062 | 0.094563 |
| 6 | -1.5 | 2000000 | 0.031623 | 1 | 0.0006 | 0.002062 | 8.2E-05 | 0.039556 |
| 7 | -1.5 | 20000000 | 0.031623 | 1 | 0.0002 | 0.00065 | 1E-05 | 0.015529 |
| 8 | -1.5 | 200000000 | 0.031623 | 1 | 6E-05 | 0.000205 | 1.2E-06 | 0.00586 |
| 3 | -1 | 2000 | 0.1 | 0.99975 | 0.0198 | 0.070712 | 0.0168 | 0.237616 |
| 4 | -1 | 20000 | 0.1 | 0.99998 | 0.0063 | 0.02127 | 0.00237 | 0.111471 |
| 5 | -1 | 200000 | 0.1 | 1 | 0.002 | 0.00658 | 0.00031 | 0.046535 |
| 6 | -1 | 2000000 | 0.1 | 1 | 0.0006 | 0.002062 | 3.8E-05 | 0.018197 |
| 7 | -1 | 20000000 | 0.1 | 1 | 0.0002 | 0.00065 | 4.4E-06 | 0.006837 |
| 8 | -1 | 200000000 | 0.1 | 1 | 6E-05 | 0.000205 | 5.1E-07 | 0.002501 |
| 3 | -0.5 | 2000 | 0.316228 | 0.99975 | 0.0198 | 0.070712 | 0.01035 | 0.146395 |
| 4 | -0.5 | 20000 | 0.316228 | 0.99998 | 0.0063 | 0.02127 | 0.00133 | 0.06264 |
| 5 | -0.5 | 200000 | 0.316228 | 1 | 0.002 | 0.00658 | 0.00016 | 0.024766 |
| 6 | -0.5 | 2000000 | 0.316228 | 1 | 0.0006 | 0.002062 | 1.9E-05 | 0.009343 |
| 7 | -0.5 | 20000000 | 0.316228 | 1 | 0.0002 | 0.00065 | 2.2E-06 | 0.003422 |
| 8 | -0.5 | 200000000 | 0.316228 | 1 | 6E-05 | 0.000205 | 2.5E-07 | 0.001228 |
| 3 | 0 | 2000 | 1 | 0.99975 | 0.0198 | 0.070712 | 0.00667 | 0.094379 |
| 4 | 0 | 20000 | 1 | 0.99998 | 0.0063 | 0.02127 | 0.00084 | 0.039495 |
| 5 | 0 | 200000 | 1 | 1 | 0.002 | 0.00658 | 0.0001 | 0.015393 |
| 6 | 0 | 2000000 | 1 | 1 | 0.0006 | 0.002062 | 1.2E-05 | 0.005748 |
| 7 | 0 | 20000000 | 1 | 1 | 0.0002 | 0.00065 | 1.4E-06 | 0.00209 |
| 8 | 0 | 200000000 | 1 | 1 | 6E-05 | 0.000205 | 1.5E-07 | 0.000746 |

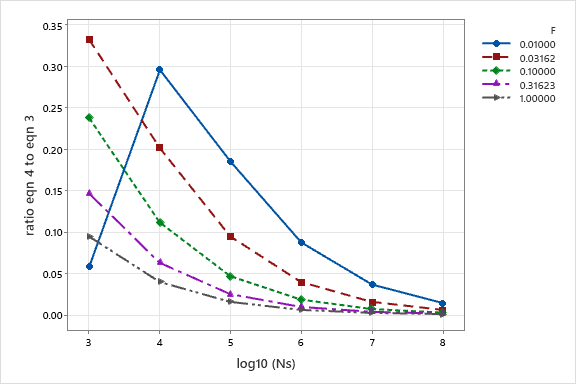

Fig. S1. Plot of the data in Table S1. Times from to fixation are always lower without random mating than with it. The y-axis shows the ratio of the time to fixation from equation 3 to that from equation 4. The aberrant point (log_10_(Ns) =3, F=0.01) results from an inadequacy of the approximations made in deriving equation 4. Note that the ratio declines as F or Ns increase, getting below 5% when Ns>100,000 and F>0.1.
