## Supplementary Material 2 for "Times birds need to evolve mating cues under allopatry and parapatry"

**Supplementary Material 2:** **Extended methods for Section 2 of Times birds need to evolve mating cues under allopatry and parapatry**

Richard M. Sibly^1*^ and Robert N. Curnow^2^

Two diploid loci are modelled, one determining local adaptation, the other a mating cue. The model of local adaptation has one locus with two alleles P and Q in an environment consisting of two niches with some migration between niches prior to mating, as depicted in Fig. 1. The locus determines ecological adaptation to one niche or the other. A large population is assumed so that the dynamics are deterministic. Generations are discrete and individuals die after mating. The life histories occur in the following order. At the start of each generation individuals in each niche mate at random, and all mating individuals obtain the same number of offspring. The number of offspring of each genotype that survive in each niche is the product of its initial frequency and its fitness. Population regulation then returns population numbers to their initial values. Finally some individuals migrate between niches, as shown in Fig. 1, leading to the start of the next generation.

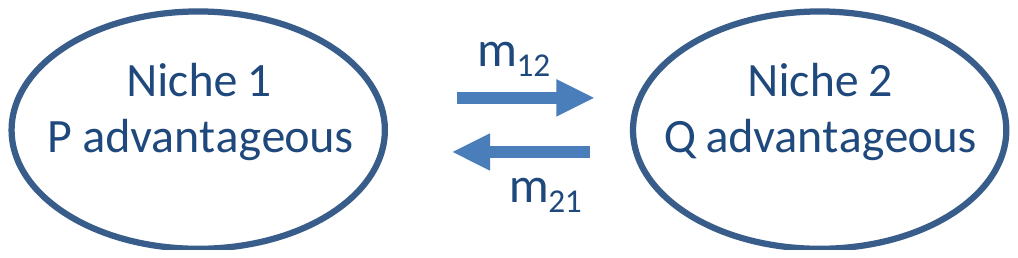

Fig. 1. Conceptual overview of the model. For clarity the niches are shown distinct, but in nature may be contiguous or overlap. m_12_ and m_21_ specify the proportion of individuals in one niche that migrate to the other each generation after viability selection and population regulation have occurred. When analysing the model it is supposed that Q is disadvantageous in niche 1 (i.e., s_1_, defined below, is negative) but advantageous in niche 2 (i.e., s_2_ is positive), while PP homozygotes have fitness 1 in both niches. There are no sex differences in fitnesses or migration rates.

| Genotype | PP | PQ | QQ |
| --- | --- | --- | --- |
| Fitnesses in our notation | 1 | 1+hs | 1+s |
| Frequency after mating | p^2^ | 2pq | q^2^ |

Table 1. Assignment of fitnesses to genotypes. h is the level of dominance of the Q allele. p and q are the frequencies of the P and Q alleles respectively; p+q=1.

The model of local adaptation shown in Fig. 1 and Table 1 consists of a single locus with three genotypes, PP, PQ and QQ. The model was extended to include a second mating cue locus, independent of and not linked to the PQ locus, with two neutral alleles C and D coding for a mating cue such as plumage colour. C is assumed dominant to D. Carriers of C are assumed to mate with other carriers of C with probability α but otherwise at random with probability (1 – α). Similarly DD individuals mate with other DD individuals with probability α but otherwise at random with probability (1 – α). There are three possible genotypes at the CD locus, so considering the two loci there are nine possible genotypes: CCQQ, CCQP, CCPP; CDQQ, CDQP, CDPP; DDQQ, DDQP and DDPP. We write their frequencies r,s,t,u,v,w,x,y and z, respectively.

We now derive the recurrence equations showing how the frequencies of the genotypes change under viability selection at the PQ locus. We begin by calculating the relative frequencies of offspring genotypes for all possible genotype crosses in an isolated niche with α=1. These frequencies are shown in Table 2. The first column gives the genotypes and frequencies of fathers, those of mothers are in the top row. The table is symmetrical.

|  | CCQQ, *r* | CCQP, *s* | CCPP, *t* | CDQQ, *u* | CDQP, *v* | CDPP, *w* | DDQQ, *x* | DDQP, *y* | DDPP, *z* |
| --- | --- | --- | --- | --- | --- | --- | --- | --- | --- |
| CCQQ, *r* | *f^2^r^2^/A* | *fgrs/A* | *frt/ A* | *f^2^ru/A* | *fgrv/A* | *frw/A* | 0 | 0 | 0 |
| CCQP, *s* | *fgrs/A* | *g^2^s^2^/A* | *gst/A* | *fgsu/A* | *g^2^sv/A* | *gsw/A* | 0 | 0 | 0 |
| CCPP, *t* | *frt/A* | *gst/A* | *t^2^/A* | *ftu/A* | *gtv/A* | *tw/A* | 0 | 0 | 0 |
| CDQQ, *u* | *f^2^ru/A* | *fgsu/A* | *ftu/A* | *f^2^u^2^/A* | *fguv/A* | *fuw/A* | 0 | 0 | 0 |
| CDQP, *v* | *fgrv/A* | *g^2^sv/A* | *gtv/A* | *fguv/A* | *g^2^v^2^/A* | *gvw/A* | 0 | 0 | 0 |
| CDPP, *w* | *frw/A* | *gsw/A* | *tw/A* | *fuw/A* | *gvw/A* | *w^2^/A* | 0 | 0 | 0 |
| DDQQ, *x* | 0 | 0 | 0 | 0 | 0 | 0 | *f^2^x^2^/B* | *fgxy/B* | *fxz/B* |
| DDQP, *y* | 0 | 0 | 0 | 0 | 0 | 0 | *fgxy/B* | *g^2^y^2^/B* | *gyz/B* |
| DDPP, *z* | 0 | 0 | 0 | 0 | 0 | 0 | *fxz/B* | *gyz/B* | *z^2^/B* |

Table 2. Relative frequencies of the genotypes of the surviving offspring of all possible genotype crosses in an isolated niche for the case α=1. A=r+s+t+u+v+w, B=x+y+z, f=1+s, g=1+hs.

The frequencies in the next generation are obtained from Table 2. In an isolated niche the frequency of offspring of the CCQQ genotype at the end of the next generation is:

$$r^{'}={\alpha(f}^{2}r^{2}+fgrs+f^{2}ru+\frac{1}{2}fgrv+\frac{1}{4}g^{2}s^{2}+\frac{1}{2}fgsu+\frac{1}{4}g^{2}sv+\frac{1}{4}f^{2}u^{2}+\frac{1}{4}fguv+\frac{1}{16}g^{2}v^{2})/A+(1-\alpha)(f^{2}r^{2}+fgrs+f^{2}ru+\frac{1}{2}fgrv+\frac{1}{4}g^{2}s^{2}+\frac{1}{2}fgsu+\frac{1}{4}g^{2}sv+\frac{1}{4}f^{2}u^{2}+\frac{1}{4}fguv+\frac{1}{16}g^{2}v^{2})$$

Similarly the frequencies of the other genotypes in the next generation are:

$$s^{'}=\alpha(fgrs+frt^{2}+\frac{1}{2}fgrv+frw+\frac{1}{2}g^{2}s^{2}+gst+\frac{1}{2}fgsu+\frac{1}{2}g^{2}sv+\frac{1}{2}gsw+ftu+\frac{1}{2}gtv+\frac{1}{4}fguv+\frac{1}{2}fuw+\frac{1}{8}g^{2}v^{2}+1/4gvw)/A+(1-\alpha)(fgrs+frt^{2}+\frac{1}{2}fgrv+frw+\frac{1}{2}g^{2}s^{2}+gst+\frac{1}{2}fgsu+\frac{1}{2}g^{2}sv+\frac{1}{2}gsw+ftu+\frac{1}{2}gtv+\frac{1}{4}fguv+\frac{1}{2}fuw+\frac{1}{8}g^{2}v^{2}+1/4gvw)$$

$$t^{'}=\alpha{(\frac{1}{4}g}^{2}s^{2}+gst+\frac{1}{4}g^{2}sv+\frac{1}{2}gsw+t^{2}+\frac{1}{2}gtv+tw+\frac{1}{16}g^{2}v^{2}+\frac{1}{4}gvw+\frac{1}{4}w^{2})/A+(1-\alpha){(\frac{1}{4}g}^{2}s^{2}+gst+\frac{1}{4}g^{2}sv+\frac{1}{2}gsw+t^{2}+\frac{1}{2}gtv+tw+\frac{1}{16}g^{2}v^{2}+\frac{1}{4}gvw+\frac{1}{4}w^{2})$$

$$u^{'}=\alpha(f^{2}ru+\frac{1}{2}fgrv+\frac{1}{2}fgsu+\frac{1}{4}g^{2}sv+\frac{1}{2}f^{2}u^{2}+\frac{1}{2}fguv+\frac{1}{8}g^{2}v^{2})/A+(1-\alpha)(f^{2}ru+\frac{1}{2}fgrv+\frac{1}{2}fgsu+\frac{1}{4}g^{2}sv+\frac{1}{2}f^{2}u^{2}+\frac{1}{2}fguv+\frac{1}{8}g^{2}v^{2}+2f^{2}rx+fgry+fgsx+\frac{1}{2}g^{2}sy+f^{2}ux+\frac{1}{2}fguy+\frac{1}{2}fgvx+\frac{1}{4}g^{2}vy)$$

$$v^{'}=\alpha(\frac{1}{4}fgrv+frw+\frac{1}{2}fgsu+\frac{1}{2}g^{2}sv+\frac{1}{2}gsw+ftu+\frac{1}{4}gtv+\frac{1}{2}fguv+fuw+\frac{1}{4}fgrv+\frac{1}{4}gtv+\frac{1}{4}g^{2}v^{2}+\frac{1}{2}gvw)/A +(1-\alpha)(\frac{1}{4}fgrv+frw+\frac{1}{2}fgsu+\frac{1}{2}g^{2}sv+\frac{1}{2}gsw+ftu+\frac{1}{4}gtv+\frac{1}{2}fguv+fuw+\frac{1}{4}fgrv+\frac{1}{4}gtv+\frac{1}{4}g^{2}v^{2}+\frac{1}{2}gvw+fgry+2frz+fgsx+g^{2}sy+gsz+2ftx+gty+\frac{1}{2}fguy+fuz+\frac{1}{2}fgvx+\frac{1}{2}g^{2}vy+\frac{1}{2}gvz +fwx+\frac{1}{2}gwy)$$

$$w^{'}=\alpha(\frac{1}{4}g^{2}sv+\frac{1}{2}gsw+\frac{1}{2}gtv+tw+\frac{1}{8}g^{2}v^{2}+\frac{1}{2}gvw+\frac{1}{2}w^{2})/A+(1-\alpha)(\frac{1}{4}g^{2}sv+\frac{1}{2}gsw+\frac{1}{2}gtv+tw+\frac{1}{8}g^{2}v^{2}+\frac{1}{2}gvw+\frac{1}{2}w^{2}+\frac{1}{2}g^{2}sy + gsz+gty+2tz+\frac{1}{4}g^{2}vy+\frac{1}{2}gvz+\frac{1}{2}gwy+2wz)$$

$$x’=\alpha(\frac{1}{4}f^{2}u^{2}+\frac{1}{4}fguv+\frac{1}{16}g^{2}v^{2})/A+\alpha(f^{2}x^{2}+fgxy+\frac{1}{4}g^{2}y^{2})/B+(1-\alpha)(\frac{1}{4}f^{2}u^{2}+\frac{1}{4}fguv+\frac{1}{16}g^{2}v^{2}+f^{2}x^{2}+fgxy+\frac{1}{4}g^{2}y^{2}+\frac{1}{2}f^{2}ux+\frac{1}{4}fguy+\frac{1}{4}fgvx+\frac{1}{8}g^{2}vy)$$

$$y’=\alpha(\frac{1}{4}fguv+\frac{1}{2}fuw+\frac{1}{8}g^{2}v^{2}+\frac{1}{4}gvw)/A+\alpha(fgxy+fxz+\frac{1}{2}g^{2}y^{2}+gyz+fxz)/B+(1-\alpha)(\frac{1}{4}fguv+\frac{1}{2}fuw+\frac{1}{8}g^{2}v^{2}+\frac{1}{4}gvw+fgxy+fxz+\frac{1}{2}g^{2}y^{2}+gyz+fxz+\frac{1}{4}fguy+\frac{1}{2}fuz+\frac{1}{4}fgvx)+\frac{1}{4}g^{2}vy+\frac{1}{4}gvz+\frac{1}{2}fwx+\frac{1}{4}gwy)$$

$$z’=\alpha(\frac{1}{16}g^{2}v^{2}+\frac{1}{4}gvw+\frac{1}{4}w^{2})/A+\alpha(\frac{1}{4}g^{2}y^{2}+gyz+z^{2})/B+(1-\alpha)(\frac{1}{16}g^{2}v^{2}+\frac{1}{4}gvw+\frac{1}{4}w^{2}+\frac{1}{4}g^{2}y^{2}+gyz+z^{2}+\frac{1}{8}g^{2}vy+\frac{1}{4}gvz+\frac{1}{4}gwy+\frac{1}{2}wz)$$

These equations apply to an isolated niche, but need modification if some individuals migrate between niches. Let the population sizes in niches 1 and 2 be M_1_ and M_2_ respectively. We use this notation to allow for later applications to finite populations but in the current infinite population model it is only the ratio M_1_/M_2_ that is important. Let the proportion of individuals in niche 1 emigrating to niche 2 each generation after viability selection be m_12_, and the proportion of those in niche 2 emigrating to niche 1 be m_21_. If the frequency of offspring of genotype i produced in niches 1 and 2 are N_1i_ and N_2i_ respectively, then after migration the number in niche 1 is M_1_N_1i_(1-m_12_) + M_2_N_2i_m_21_. An analogous formula applies to niche 2, and the number after migration is M_2_N_2i_(1-m_21_) + M_1_N_1i_m_12_. Assuming the numbers moving in the two directions are equal, then M_1_m_12_ = M_2_m_21_.

This model was run by computer simulation. Further details including the computer code used to run the simulations is available with explanatory annotations in (Sibly and Curnow, 2022).

Sibly, R. M., Curnow, R. N., 2022. Sexual imprinting leads to speciation in locally adapted populations. Ecology and Evolution, Vol. 12.
